# The histone demethylase KDM5 is required for synaptic structure and function at the *Drosophila* neuromuscular junction

**DOI:** 10.1101/2020.10.12.335711

**Authors:** Helen M. Belalcazar, Emily L. Hendricks, Sumaira Zamurrad, Faith L.W. Liebl, Julie Secombe

## Abstract

Mutations in the genes encoding the KDM5 family of histone demethylases are observed in individuals with intellectual disability (ID). Despite clear evidence linking KDM5 function to neurodevelopmental pathways, how this family of proteins impacts transcriptional programs to mediate synaptic structure and activity remains unclear. Using the *Drosophila* larval neuromuscular junction (NMJ), we show that KDM5 is required for neuroanatomical development and synaptic function. The JmjC-domain encoded histone demethylase activity of KDM5, which is expected to be diminished by many ID-associated alleles and required for appropriate synaptic morphology and neurotransmission. The C5HC2 zinc finger of KDM5 is also involved, as an ID-associated mutation in this motif reduces NMJ bouton number but increases bouton size. KDM5 therefore uses demethylase-dependent and independent mechanisms to regulate NMJ structure and activity, highlighting the complex nature by which this chromatin modifier carries out its neuronal gene regulatory programs.

## Introduction

In the central nervous system (CNS), misregulation of gene expression has profound effects on cognitive and other neurological functions (Ronan et al., 2013). The recent expansion of genomic analyses in patients with neurodevelopmental disorders (NDDs) has dramatically increased our understanding of the genetic contributors to cognitive impairment. These studies revealed an association between mutations in genes encoding <u>lysine</u> <u>d</u>emethylase <u>5</u> (KDM5) family proteins and NDDs, including intellectual disability (ID) (Vallianatos and Iwase, 2015). KDM5 proteins are chromatin modifiers that enzymatically remove di- and/or trimethylated lysine 4 of histone H3 (H3K4me2/3) through activity of a conserved Jumonji C (JmjC) domain (Christensen et al., 2007; Iwase et al., 2007; Secombe et al., 2007; Yamane et al., 2007). The H3K4me2/3 chromatin marks are predominantly found at the promoter region of transcriptionally active genes and changes to these covalent histone modifications can impact transcriptional consistency (Barski et al., 2007; Benayoun et al., 2014).

There are four *KDM5* paralogs in humans. *KDM5A* and *KDM5B* are autosomal while *KDM5C* and *KDM5D* are X- and Y-linked, respectively. *KDM5A*, *KDM5B*, and *KDM5C* are each expressed broadly in the brain and 106 distinct mutations in these genes have been found in individuals with NDDs (Carmignac et al., 2020; Stenson et al., 2017). Clinically, mutations in *KDM5C* are primarily associated with syndromic ID (OMIM#300534), with patients presenting with low cognitive function and comorbid features such as autism, aggressive behavior, spastic paraplegia and microcephaly (Abidi et al., 2008; Adegbola et al., 2008; Carmignac et al., 2020; Goncalves et al., 2014; Jensen et al., 2005; Rujirabanjerd et al., 2010; Santos-Reboucas et al., 2011). Many of these cognitive and motor deficits are recapitulated in *Kdm5c* knockout mice, consistent with the evolutionary conservation of *KDM5C’s* neuronal functions (Iwase et al., 2016; Scandaglia et al., 2017). Genome-wide studies of patients with ID and their families have also revealed a link between mutations in *KDM5B* and autosomal recessive cognitive impairment (Al-Mubarak et al., 2017; Faundes et al., 2018; Lebrun et al., 2018) (OMIM#618109). While examination of *Kdm5b* knockout mice firmly establishes a direct genetic link between loss of *KDM5B* and ID, the range and severity of patient phenotypes associated with these mutations are less defined than those associated with *KDM5C*-induced ID (Faundes et al., 2018; Martin et al., 2018). Mutations in *KDM5A* are infrequently observed in individuals with NDDs, with only two mutations described to date; one in a patient with autism and the other in a patient with ID (Butler et al., 2015; Najmabadi et al., 2011). Because studies of *Kdm5a* knockout mice have not examined changes in cognition (Klose et al., 2007), the extent to which loss of KDM5A contributes to patient phenotypes remains unclear.

Approximately half of the 106 reported *KDM5A*, *KDM5B* and *KDM5C* NDDs patient mutations are predicted to reduce or eliminate protein function due to the introduction of a stop codon or frame shift (Brookes et al., 2015; Goncalves et al., 2014; Najmabadi et al., 2011). The remaining genetic changes observed in NDDs patients are missense mutations and are best characterized for *KDM5C*. Several missense mutations reduce protein stability and are likely to mimic loss of KDM5C (Brookes et al., 2015). However, at least three ID patient-associated missense mutations do not alter overall protein levels (Peng et al., 2015; Vallianatos et al., 2018), suggesting that they could affect a subset of KDM5A or KDM5C functions that are particularly important for neuronal morphology and/or activity. Analyses of missense alleles, therefore, provides an opportunity to understand the neuronal functions of KDM5 proteins.

The prevailing model of *KDM5*-induced ID suggests that all mutations diminish demethylase activity, which is critical for neuronal development and function. In support of this, *in vitro* studies using recombinant KDM5C harboring ID-associated missense alleles within or outside the enzymatic JmjC domain reduce activity toward a histone peptide substrate up to 2-fold (Iwase et al., 2007; Rujirabanjerd et al., 2010; Tahiliani et al., 2007). Additionally, *in vivo* analyses show that the cognitive deficits observed in *Kdm5c* knockout mice can be partially ameliorated by genetically lowering levels of the histone methyltransferase KMT2A (*Kmt2a* heterozygotes). Consistent with these observations, a *Drosophila* mutant specifically lacking KDM5 demethylase activity show learning and memory deficits (Zamurrad et al., 2018). Contrasting these findings, it should be noted that some ID-associated missense mutations in *KDM5C* do not show detectable deficits to *in vitro* demethylase activity (Tahiliani et al., 2007; Vallianatos et al., 2018). This suggests that non-enzymatic mechanisms of gene regulation by KDM5C may also be important for its neural functions. Consistent with this, mammalian KDM5A, KDM5B and *Drosophila* KDM5 can regulate gene expression independent of their demethylase activity, although the molecular mechanisms and physiological importance of these non-canonical activities remain obscure (Cao et al., 2014; Liu and Secombe, 2015; Zou et al., 2014). KDM5 family proteins may therefore use different domains to transcriptionally regulate distinct sets of target genes that are important for proper brain development and function.

There is still much to be learned regarding the cellular consequences of mutations in *KDM5* family genes. Structural changes in neuronal morphology and circuitry are often observed in neurodevelopmental and neuropsychiatric disorders (Forrest et al., 2018). For example, analyses of post mortem brain samples from individuals with ID show atypical dendritic arborization and spine morphology (Bagni and Zukin, 2019; Nishiyama, 2019). This phenotype has yet to be examined in patients with mutations in *KDM5* family genes. However, knockdown of *Kdm5c* in cultured rat cerebellar granular neurons results in changes in dendritic arborization (Iwase et al., 2007) and *Kdm5c* knockout mice show altered cortical neuron spine density (Iwase et al., 2016; Scandaglia et al., 2017). While dendritic spine morphology can be altered by neuronal activity (Nagerl et al., 2004; Verpelli et al., 2010), the extent to which synaptic function is disrupted by mutations in *KDM5* genes remains uncharacterized. Additionally, the *Kdm5c* gene expression program that mediates the morphological changes in mouse and rat cells and whether it relies on the demethylase activity of KDM5C, remains largely unexplored.

The glutamatergic *Drosophila* neuromuscular junction (NMJ) serves as a well-established model for addressing questions of neural development, morphology, and synaptic function (Li et al., 2007; Rasse et al., 2005; Roos et al., 2000). The connection between motor neurons and muscle contains proteins homologous to those found in neuronal mammalian CNS excitatory synapses (Koh et al., 2000). Mutations in the human orthologs of genes encoding synaptic proteins, including ionotropic and metabotropic glutamate receptors, the scaffold protein PSD95/Dlg, the phosphoprotein Synapsin and the cell-adhesion proteins Neurexin and Neuroligin, lead to altered cognition (Moretto et al., 2018; Volk et al., 2015). *Drosophila* NMJs are established during the embryonic stage of development when motor neuron axons contact the muscle and form branches consisting of synaptic boutons arranged in a stereotypic “beads on a string” morphology (Gramates and Budnik, 1999). The addition of new boutons is required during larval growth to maintain synaptic strength (Menon et al., 2013) and occurs by the budding or division of mature boutons or by *de novo* formation (Zito et al., 1999). Analyses of the pathways regulating synaptic bouton quantity or morphology have elucidated important neuropathological mechanisms for ID-associated genes (Bellosta and Soldano, 2019; Tian et al., 2017).

*Drosophila kdm5*, also known as *little imaginal discs* (*lid*), is ubiquitously expressed and essential for developmental timing and adult survival (Drelon et al., 2018; Drelon et al., 2019; Secombe et al., 2007). As *Drosophila* KDM5 shares homology with all four mammalian KDM5 proteins, analysis of its function is expected to reveal activities relevant to *KDM5A*, *KDM5B* and *KDM5C*-induced ID. In this study, we show that KDM5 is required in motor neurons for proper NMJ morphology and function. Specifically, we demonstrate that the histone demethylase activity of KDM5 is a key contributor to its regulatory effects at the synapse as animals lacking catalytic activity (*kdm5^JmjC*^*) show a reduction in the number of type Ib synaptic boutons and a decrease in evoked glutamate release. Additionally, we find that KDM5 requires the C5HC2 zinc finger, found immediately adjacent to the enzymatic JmjC domain, to perform its neuroregulatory functions. A missense allele equivalent to a patient-associated mutation in *KDM5C*, *kdm5^L854F^*, which is within the C5HC2 motif, reduces the number of synaptic boutons but increases their size. Interestingly, this ID-associated allele does not appear to dramatically alter histone demethylase activity, suggesting that KDM5 uses canonical and non-canonical mechanisms of gene regulation to properly form and maintain NMJs. Consistent with these data, RNA-seq analyses of the ventral nerve cord (VNC), where the nuclei of motor neurons reside, revealed that mutations in either the JmjC (*kdm5^JmjC*^)* or C5HC2 (*kdm5^L854F^*) domain cause distinct transcriptional changes. Combined, these data advance our understanding of the neuronal activities of KDM5 and elucidate the functions of discrete domains that mediate its gene regulatory programs in neurons.

## Results

### KDM5 is required presynaptically for proper NMJ structure

To assess the role of KDM5 at the synapse, we examined the NMJ structure in *kdm5^140^* null mutant animals. We immunolabeled third instar larval NMJs with the neuronal membrane marker horseradish peroxidase (HRP) and the postsynaptic scaffold protein Discs large (Dlg), which is the *Drosophila* homolog of PSD-95. Based on this staining, we quantified the number and size of type Ib synaptic boutons at muscle 4 of the third abdominal segment, which are predominantly responsible for muscle contraction (Newman et al., 2017), in *kdm5^140^* null mutants. Similar to our previous studies, we matched animals developmentally rather than chronologically due to the developmental delay of *kdm5^140^* larvae (Drelon et al., 2018; Drelon et al., 2019). In addition, we normalized the number of boutons to muscle size to account for any slight differences in larval size. These analyses revealed that loss of *kdm5* alters NMJ morphology by decreasing the number and increasing the size of synaptic boutons (Figure 1A - B, F, H).

**Figure 1.**
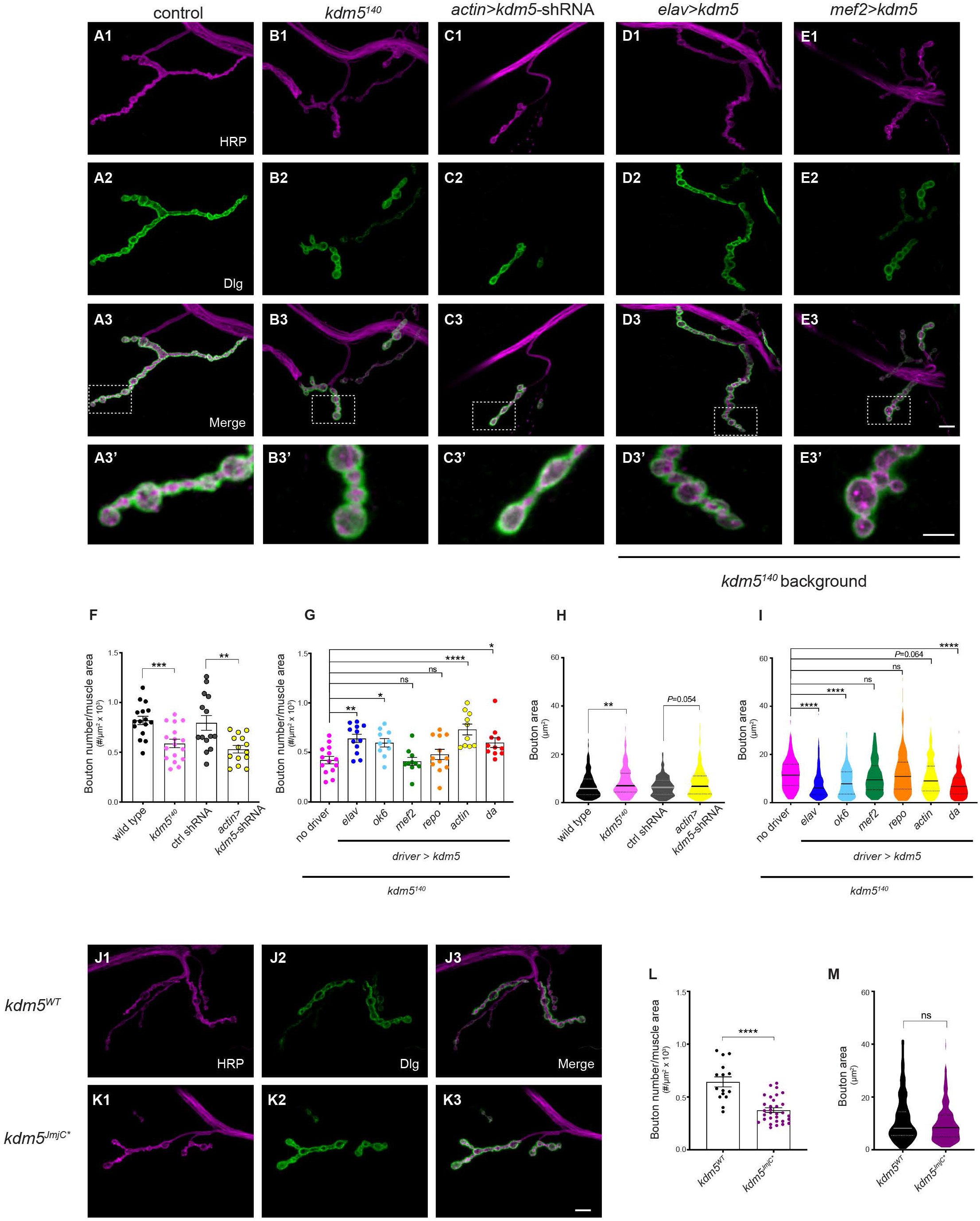
*kdm5* expression is required in motor neurons for normal growth of the larval NMJ. **A1-A3** NMJ morphology at muscle 4 of the abdominal segment 3 (NMJ4-A3) of control strain that is wild type for *kdm5* (*w^1118^*) third instar larvae labeled with the presynaptic marker HRP (magenta; A1), the postsynaptic marker Dlg (green; A2) and the merge (A3). (**B1-3**) *kdm5^140^* homozygous larva NMJs revealing decrease in number and increase in size of type Ib synaptic boutons compared to wild type. (**C1-3**) NMJ from *kdm5* knockdown larva (actin>*kdm5*-shRNA) showing similar phenotype to *kdm5^140^*. (**D1-3**) NMJ from *kdm5^140^* larva, presynaptically re-expressing *kdm5* (*kdm5^140^*; *elav>UAS-kdm5*), showing rescue of the null phenotype. (**E1-3**) NMJ from a *kdm5^140^* larva re-expressing *kdm5* in muscle tissue (*kdm5^140^*; *mef2>UAS-kdm5*). (**A3’-E3’**) Boxed areas in A3-E3 showing higher magnification of type Ib boutons. Scale bars: 10μm (E3) and 5μm (E3’). **F**. Quantification of type Ib bouton number normalized to muscle surface area in *kdm5* loss of function and knockdown of the indicated genotypes. \*\*\**P*=0.0001 (Mann-Whitney test), \*\**P*=0.0041 (Unpaired t test). *w^1118^* n=20, *kdm5^140^* n=25, *actin>kdm5;shRNA* n=14, control (kdm5;shRNA) n=14. **G**. Quantification of type Ib bouton number normalized to muscle surface area in rescue experiments of indicated genotypes. \*\*\**P*<0.0001, \*\**P*=0.0029, \**P*=0.0036 (*OK6*), \**P*=0.0282 (*da*) ns = not significant (*mef2, repo*) (one-way ANOVA Dunnett’s multiple comparisons). No driver-*kdm5^140^*; *UAS.kdm5* n=14, *elav>kdm5* n=12, *OK6*>*kdm5* n=10, *mef2>kdm5* n=10, *repo>kdm5* n=12, *actin>kdm5* n=10, *da*>*kdm5* n=11. **H**. Quantification of type Ib bouton size in *kdm5^140^* larvae and ubiquitous knockdown (*actin>kdm5-shRNA*). Violin plots show the frequency distribution of bouton surface area indicating the median and quartiles. \*\**P*=0.0020 (Mann-Whitney test). *w^1118^*n=327, *kdm5^140^* n=280, *actin>kdm5;shRNA* n=198, control (*kdm5*;shRNA) n=281. **I**. Quantification of type Ib bouton size in rescue experiments using the genotypes shown. Violin plots show the frequency distribution of bouton surface area indicating the median and quartiles \*\*\*\**P*<0.0001, *P*=0.064 ns = not significant (*mef2, repo*) (Kruskal-Wallis Dunn’s multiple comparisons). No driver-*kdm5^140^*; *UAS.kdm5* n=149, *elav>kdm5* n=245, *OK6>kdm5* n=191, *mef2>kdm5* n=123, *repo>kdm5* n=254, *actin>kdm5* n=181, *da>kdm5* n=201. **J1-K3**. NMJ4-A3 of wild-type (*kdm5^WT^*; J1-3) and demethylase inactive (*kdm5^JmjC*^*; K1-3) third instar larva stained with HRP (J1, K1), Dlg (J2, K2), and the merged panels (J3, K3) showing that loss of enzymatic activity leads to decrease in bouton number but does not affect bouton size. Scale bar: 10μm. **L**. Quantification of type Ib bouton number normalized to muscle surface area in *kdm5^WT^* and *kdm5^JmjC*^*. \*\*\*\**P*<0.0001 (Unpaired t test). *kdm5^WT^* n=15, *kdm5^JmjC*^* n=31. **M.** Quantification of type Ib bouton size in *kdm5^WT^* and *kdm5^JmjC*^*. Violin plots show the frequency distribution of bouton surface area indicating the median and quartiles. ns = not significant (Mann-Whitney test). *kdm5^WT^* n=234, *kdm5^JmjC*^* n=388. Bars represent mean with SEM.

To independently validate this phenotype, we used a well-established inducible *kdm5* short hairpin RNA (shRNA) transgene that reduces KDM5 protein levels by ~75% (Chen et al., 2019; Liu and Secombe, 2015) (Figure S1). Ubiquitous knock down of *kdm5* using *Actin5c-Gal4* recapitulated the reduced bouton number phenotype that we observed in *kdm5^140^* larvae (Figure 1C, F, H). *kdm5* knock down larvae also showed a modest increase in NMJ bouton size (P=0.05) that was less drastic, but consistent with the *kdm5^140^* null phenotype. To rule out any effects of an altered larval growth rate on the observed NMJ phenotypes, we examined the developmental timing of *Actin5c>kdm5-shRNA* larvae and found that larval growth was not delayed in the same manner as the null allele (Figure S1). This suggests that KDM5 acts through distinct mechanisms to regulate larval growth rate and NMJ morphology. Combined, our analyses of *kdm5* null mutant and knockdown larvae show that KDM5 is required to promote NMJ bouton number and to restrict bouton size.

To determine where KDM5 function is required to mediate its effects on NMJ morphology, we restored *kdm5* expression in specific cell types within *kdm5^140^* protein-null animals. To achieve this, we used a *UAS-kdm5* transgene that drives low levels of expression (~2-fold increase over endogenous levels) and a range of *Gal4* drivers (Drelon et al., 2019; Li et al., 2010; Secombe et al., 2007). Re-expression of *kdm5* ubiquitously (*Actin5c*-*Gal4* and *da*-*Gal4*), pan-neuronally (*elav*-*Gal4*), or in motor neurons (*OK6*-*Gal4*) significantly rescued the bouton number deficit of *kdm5^140^* larvae (Figure 1D, G). Restoring *kdm5* expression in muscles (*Mef2*-*Gal4*) or glia (*repo*-*Gal4*), however, did not (Figure 1E, G). Similar findings were observed with respect to the increased bouton size phenotype in *kdm5^140^* larvae. Neuronal re-expression of *kdm5* using *elav*-*Gal4* and *OK6*-*Gal4* restored bouton size similar as wild type, whereas muscle or glial expression did not (*Mef2*-*Gal4* and *repo*-*Gal4*, respectively) (Figure 1I). These data suggest that KDM5 impacts NMJ growth by regulating presynaptic gene expression.

A majority of ID-associated mutations in *KDM5* family genes are expected to compromise histone demethylase activity, which has been proposed to play a key role in the cognitive impairment observed in patients (Vallianatos and Iwase, 2015). We therefore examined the contribution of this chromatin-modifying activity to NMJ morphology using a fly strain with two point mutations in the JmjC domain (*kdm5^JmjC*^*) that abolish enzymatic function (Drelon et al., 2018; Navarro-Costa et al., 2016; Zamurrad et al., 2018). *kdm5^JmjC*^* homozygous larvae exhibited a decrease in bouton number but no change in bouton size compared to the isogenic control strain for this mutation (*kdm5^WT^*) (Figure 1J-M). Thus, the catalytic activity of KDM5 is required for only one of the phenotypes observed in the null mutant suggesting that non-enzymatic activities are also required for proper NMJ structure.

### Characterization of fly strains harboring ID-associated mutations in the C5HC2 zinc finger domain of KDM5

Approximately 25% of *KDM5C* missense mutations found in patients with ID occur within the uncharacterized C5HC2 zinc finger motif (Human Gene Mutation Database (HGMD^®^), (Stenson et al., 2017), suggesting a key functional role for this domain in neurons. We generated three fly strains harboring ID-associated mutations in the C5HC2 domain: *kdm5^L854F^, kdm5^R873W^* and *kdm5^Y874C^*, which are equivalent to *KDM5C^L731F^*, *KDM5C^R750W^* and *KDM5C^Y751C^*, respectively (Figure 2A) (Jensen et al., 2005; Tzschach et al., 2006). These alleles were chosen because they result in moderate to severe intellectual disability and therefore may more dramatically change gene expression in neuronal lineages. It is possible that these alleles primarily affect the enzymatic activity of the adjacent JmjC domain. Indeed, *in vitro* histone demethylase assays using recombinant KDM5C revealed that mutations equivalent to KDM5^Y874C^ and KDM5^L854F^ caused a 2-fold decrease in enzymatic activity (Iwase et al., 2007). However, because the effect of these mutations on *in vivo* enzymatic activity against a nucleosomal substrate remains unknown, it is also plausible that the C5HC2 domain mediates transcriptional activities through non-enzymatic mechanisms. Similar to our approach for generating the *kdm5^JmjC*^* allele, we used a genomic rescue strategy to generate ID-associated alleles (Drelon et al., 2018; Li et al., 2010; Liu and Secombe, 2015; Navarro-Costa et al., 2016; Zamurrad et al., 2018). This involved generating fly strains lacking the endogenous *kdm5* locus and inserting the entire 11kb *kdm5* genomic region containing the point mutation and three tandem copies of the HA epitope tag into a new location on the 3^rd^ chromosome (the *attP* site at cytological location 86F). The wild-type control strain for these alleles is homozygous for the *kdm5^140^* mutation and the HA-tagged wild-type locus at 86F (*kdm5^WT^*) (Drelon et al., 2018; Zamurrad et al., 2018).

**Figure 2.**
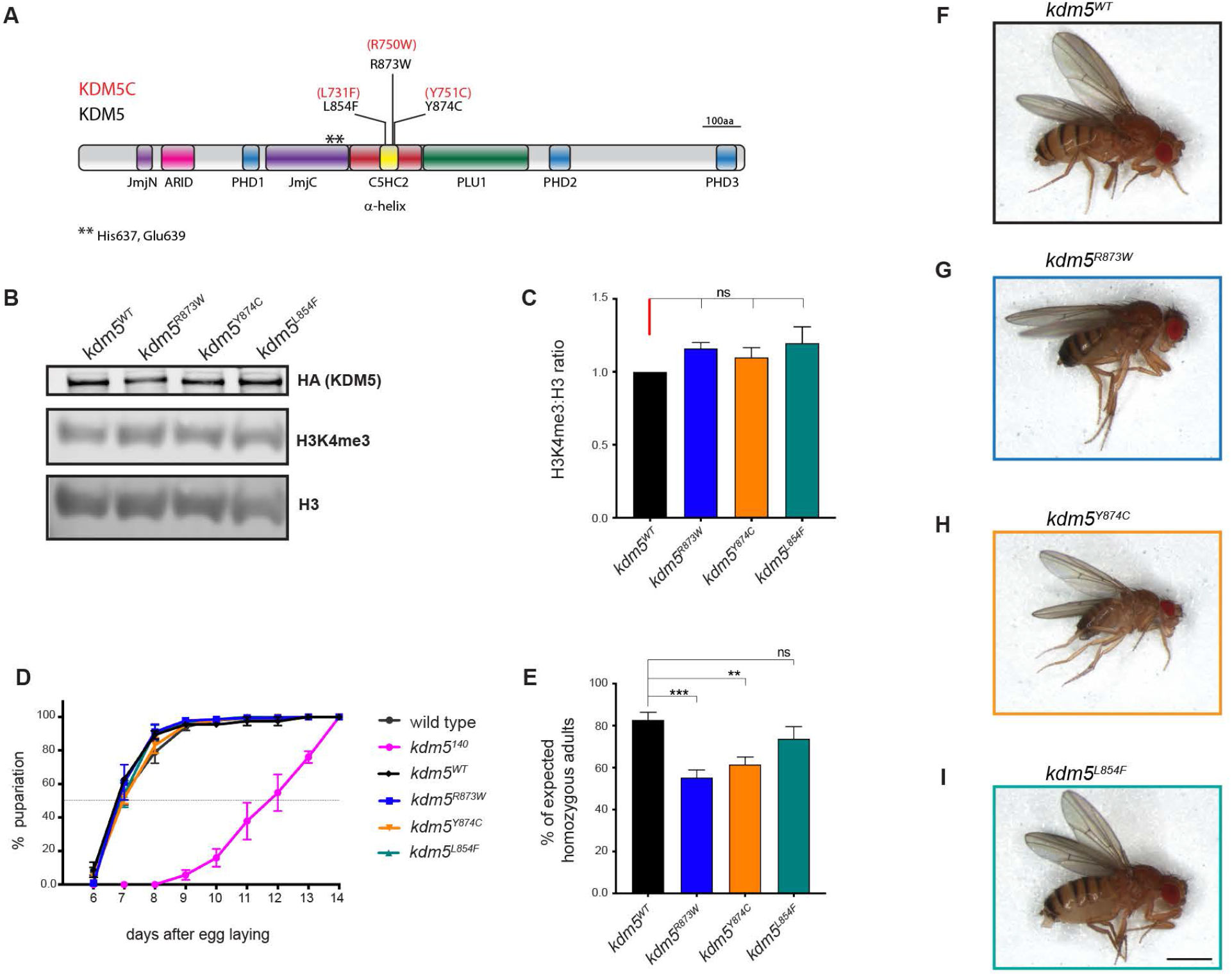
Fly strains harboring ID-associated mutations are adult viable and developmentally similar to controls. **A**. Schematic representation of *Drosophila* KDM5 showing the equivalent ID-associated mutation in human KDM5C. Stars represent the position of the two missense mutations in the JmjC domain that abolish demethylase activity in the *kdm5^JmjC*^* strain. **B**. Representative western blot using dissected larval CNS from *kdm5^WT^*, *kdm5^R873W^*, *kdm5^Y874C^* and *kdm5^L854F^* showing expression of HA-tagged KDM5 (top; 200kDa), H3K4me3 (middle) and histone H3 (bottom). **C**. Quantification H3K4me3 levels relative to total histone H3 in *kdm5^R873W^*, *kdm5^Y874C^* and *kdm5^L854F^* compared to the ratio observed in *kdm5^WT^*. There was no statistical difference in H3K4me3:H3 ratio for any mutant. ns = not significant (Kruskal-Wallis test). Results are from three different western blots. **D**. Developmental timing of the control strain *w^1118^*, *kdm5^WT^*, *kdm5^140^*, *kdm5^R873W^*, *kdm5^Y874C^*, and *kdm5^L854F^* by determining the day when 50% of the animals reach pupariation. ID mutants have the same developmental timing as *kdm5^WT^*. Total pupae quantified in three different experiments. *w^1118^* (n=88), *kdm5^140^* (n=52), *kdm5^WT^* (n=398), *kdm5^R873W^* (n=216), *kdm5^Y874C^* (n=294), *kdm5^L854F^* (n=137). **E**. Adult survival of homozygous *kdm5^WT^*, *kdm5^R873W^*, *kdm5^Y874C^*, and *kdm5^L854^* adult flies from intercrossed heterozygous parents. Data are shown as the percentage of the number of homozygous adults expected based on Mendelian expectations. \*\*\**P*= 0.0001, \*\**P*=0.0057 ns = not significant Fisher’s exact test. Total scored flies. *kdm5^WT^* (n=359), *kdm5^R873W^* (n=1164), *kdm5^Y874C^* (n=1193), *kdm5^L854F^* (n=544). **F-I**. Female flies wild type (*kdm5^WT^*) (F), *kdm5^R873W^*(G), *kdm5^Y874C^* (H) and *kdm5^L854F^* (I) showing normal size and external morphology. Scale bar: 100μm. Error bars represent SEM.

To determine whether the ID-associated alleles altered KDM5 expression, we performed an anti-HA western blot and found no change to CNS protein levels in *kdm5^L854F^, kdm5^R873W^* or *kdm5^Y874C^* animals compared to *kdm5^WT^* (Figure 2B). Because a key feature of *kdm5^140^* null mutant larvae is extended larval development that delays pupariation, we quantified the developmental timing of *kdm5^L854F^, kdm5^R873W^* and *kdm5^Y874C^* animals. Similar to our previous analyses of homozygous *kdm5^JmjC*^* animals (Drelon et al., 2018), ID mutant strains showed an identical developmental profile to *kdm5^WT^* (Figure 2D). This allowed us to examine the NMJ phenotypes of these animals without the complication of delayed larval growth rate. All three ID-associated mutant fly strains were homozygous viable and visibly morphologically normal (Figure 2E-I). *kdm5^R873W^* and *kdm5^Y874C^* adult flies occurred, however, slightly less frequently than expected. These alleles, therefore, do not dramatically alter the developmental functions of KDM5 that are required for adult viability.

Alleles of *kdm5* that abolish enzymatic activity cause a global 2-fold increase in the ratio of H3K4me3 to total histone H3, as assessed by western blot (Drelon et al., 2018; Navarro-Costa et al., 2016; Secombe et al., 2007; Zamurrad et al., 2018). To examine the effect of the KDM5^L854F^, KDM5^R873W^ and KDM5^Y874C^ mutations on histone demethylase activity *in vivo*, we similarly quantified levels of H3K4me3 using dissected larval CNS tissue. In contrast to *kdm5^JmjC*^*, the ID-associated mutants *kdm5^L854F^*, *kdm5^R873W^* and *kdm5^Y874C^* did not significantly affect H3K4me3 levels compared to total histone H3 (Figure 2B-C). While it remains possible that these mutations have modest or gene-specific effects that are undetectable by western blot, these alleles do not abolish demethylase activity *in vivo*.

To further characterize these ID-associated mutations, we examined their thermodynamic effect on KDM5 using the web server DynaMut, which calculates the change in folding Gibbs free energy (ΔΔG) by combining normal mode analysis and graph-based signatures. We also used DynaMut to measure the vibrational entropy (ΔΔSVib), a measure of molecule flexibility (Rodrigues et al., 2018). Because there is no crystal structure of *Drosophila* KDM5, we analyzed the KDM5A structure PDB:5CEH, which covers a broader N-terminal region and C-terminal helical zinc-binding domain compared to the structures available for KDM5C (Vinogradova et al., 2016) (Figure S2A). All residues investigated are conserved between *Drosophila* KDM5, human KDM5A and human KDM5C. All three mutations were predicted to thermodynamically stabilize the protein with the maximum effect shown by Y720C (Y874C in *Drosophila*) (ΔΔG: 1.759 kcal/mol), followed by L700F (L854F in *Drosophila*) (ΔΔG: 1.624 kcal/mol) (Figure S2B-D). The latter is predicted to generate new hydrophobic contacts with residues localized in the adjacent helix and increase hydrogen bonds with a residue inside the first zinc-binding site (Figure S2B). DynaMut also predicted the greatest decrease in molecule flexibility for L700F/L854F (ΔΔSVib ENCoM: −5.152 kcal.mol-1.K-1). These results, together with the position of the L700/L854 residue in the middle of the first zinc-binding site, suggests that this amino acid change may have the most dramatic impact on KDM5 function compared to the other amino acid substitutions (Figure S2A).

### The JmjC domain and the C5HC2 motif of KDM5 are required for NMJ structure and function

To determine the functional impact of ID-associated mutations in the C5HC2 domain of KDM5, we examined NMJ morphology in *kdm5^L854F^*, *kdm5^R873W^* and kdm5^Y874C^ homozygous mutant larvae. Quantification of type Ib synaptic bouton number and size showed that *kdm5^L854F^* larvae had a significantly fewer muscle 4 synaptic boutons compared to *kdm5^WT^*. In contrast, *kdm5^Y874C^* animals exhibited a mild reduction that did not meet statistical significance (*P*=0.077) while *kdm5^R873W^* bouton number was indistinguishable from *kdm5^WT^* animals (Figure 3A-E). We additionally found an increase in *kdm5^L854F^* bouton size but not in *kdm5^R873W^* or *kdm5^Y874C^* mutants when compared to *kdm5^WT^* (Figure 3F). The *kdm5^L854F^* allele therefore recapitulated both the decreased bouton number and increased bouton size phenotypes observed in the *kdm5^140^* null strain. Notably, although both *kdm5^JmjC*^* and *kdm5^L854F^* mutants show similar phenotypes with respect to bouton number, only *kdm5^JmjC*^* increases the global H3K4me3:H3 ratio (Figure 2C) (Zamurrad et al., 2018). This suggests that the enzymatic JmjC domain and the C5HC2 zinc finger of KDM5 are both required for maintaining proper NMJ morphology. Because both *kdm5^JmjC*^* and *kdm5^L854F^* adversely affected NMJ morphology without altering larval growth rate or viability, we focused the remainder of our analyses on these two alleles.

**Figure 3.**
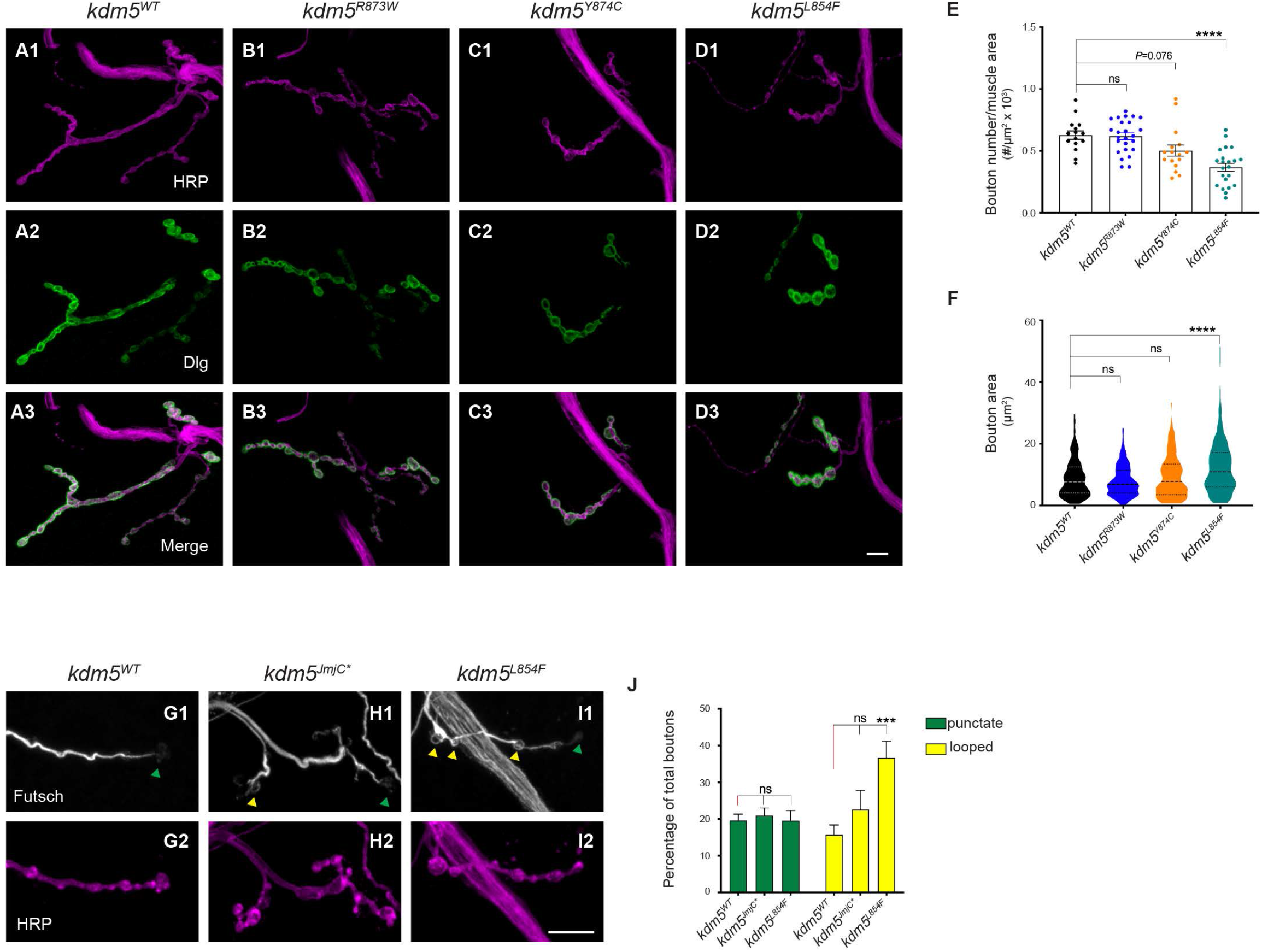
The L854F mutation in the C5HC2 domain of KDM5 phenocopies loss of KDM5 at the NMJ. **A1-D3**. NMJ morphology at muscle 4-A3 of third instar larvae labeled with Dlg (postsynaptic-green; A1, B2, C1, D1) and HRP (presynaptic-magenta; A2, B2, C2, D2). (**A1-3**) NMJ from *kdm5^WT^* larva. (**B1-3**) *kdm5^R873W^*. (**C1-C3**) *kdm5^Y874C^*. (**D1-3**) *kdm5^L854F^*. **E**. Quantification of type Ib bouton number normalized to muscle surface area from *kdm5^WT^*, *kdm5^R873W^*, *kdm5^Y874C^*, and *kdm5^L854^* larvae. \*\*\*\**P*<0.0001, ns = not significant (Kruskal-Wallis Dunn’s multiple comparisons). *kdm5^WT^* n=15, *kdm5^R873W^* n=25, *kdm5^Y874C^* n=16, *kdm5^L854F^* n=22. **F.** Quantification of type Ib bouton size from *kdm5^WT^*, *kdm5^R873W^*, *kdm5^Y874C^*, and *kdm5^L854^* larvae. Violin plots show the frequency distribution of bouton surface area indicating the median and quartiles. \*\*\**P*=0.0005, ns = not significant (Kruskal-Wallis Dunn’s multiple comparisons). *kdm5^WT^* n=309, *kdm5^R873W^* n=214, *kdm5^Y874C^* n=97, *kdm5^L854F^* n=176. **G-I**. Futsch (white) immunolabeling in segments of NMJ4-A3 showing microtubules within synaptic boutons and HRP (magenta) from *kdm5^WT^* (G1, G2) *kdm5^JmjC*^* (H1, H2) and *kdm5^L854^* (I1, I2) larvae. Green triangles indicate boutons with punctate signal and yellow triangles indicate looped boutons. **J.** Quantification of unbundled boutons, punctate (green) and looped (yellow) as percentage of total boutons in NMJ4-A3 of *kdm5^WT^*, *kdm5^JmjC*^* and *kdm5^L854^* larvae. Looped boutons were increased in *kdm5^L854F^* compared to *kdm5^WT^*. \*\*\**P*=0.0007, ns = not significant (one-way ANOVA Dunnett’s multiple comparisons). *kdm5^WT^* n=19, *kdm5^JmjC*^*=12, *kdm5^L854F^* n=14. Bars represent mean with SEM. Scale bar: 10μm.

Changes in bouton size correlate with changes in microtubule stability, which can be visualized by examining the distribution of the microtubule binding protein Futsch (Nechipurenko and Broihier, 2012; Viquez et al., 2006). We therefore stained the NMJs of *kdm5^JmjC*^* and *kdm5^L854F^* animals with anti-Futsch and quantified the number of boutons containing unbundled microtubules, which are characterized as a looped or punctuate signal (Packard et al., 2002).

There was no difference in the proportion of punctate boutons for either mutant strain. However, *kdm5^L854F^* mutant larvae showed a higher proportion of looped microtubules in boutons (36.6%) compared to *kdm5^WT^*, while *kdm5^JmjC*^* larvae did not (22.6% vs 15.7% *kdm5^WT^*) (Figure 3G-J). Thus, the increased proportion of unbundled microtubules caused by the *kdm5^L854F^* mutation may be contributing to the increase in bouton size observed in these animals. The C5HC2 domain of KDM5 may therefore be necessary for regulating microtubule dynamics at the synapse.

After establishing that mutations in *kdm5* impact NMJ morphology, we next asked whether the structural defects observed in *kdm5^JmjC*^* and *kdm5^L854F^* were associated with larval movement deficits (Kashima et al., 2017; Li et al., 2007). *kdm5^JmjC*^* and *kdm5^L854F^* third instar larvae exhibited a significant decrease in locomotor performance as indicated by quantification of crawling behavior. Specifically, both genotypes traveled a shorter total distance (path length) compared to *kdm5^WT^* (Figure 4A, E). This is likely due to reduced locomotor speed as *kdm5^JmjC*^* and *kdm5^L854F^* animals showed lower average and maximal velocities as well as average speed normalized to larval size (Figure 4B-D). Loss of KDM5 catalytic activity and the ID-associated mutation L854F in the C5HC2 domain therefore affect functional motor outputs.

**Figure 4.**
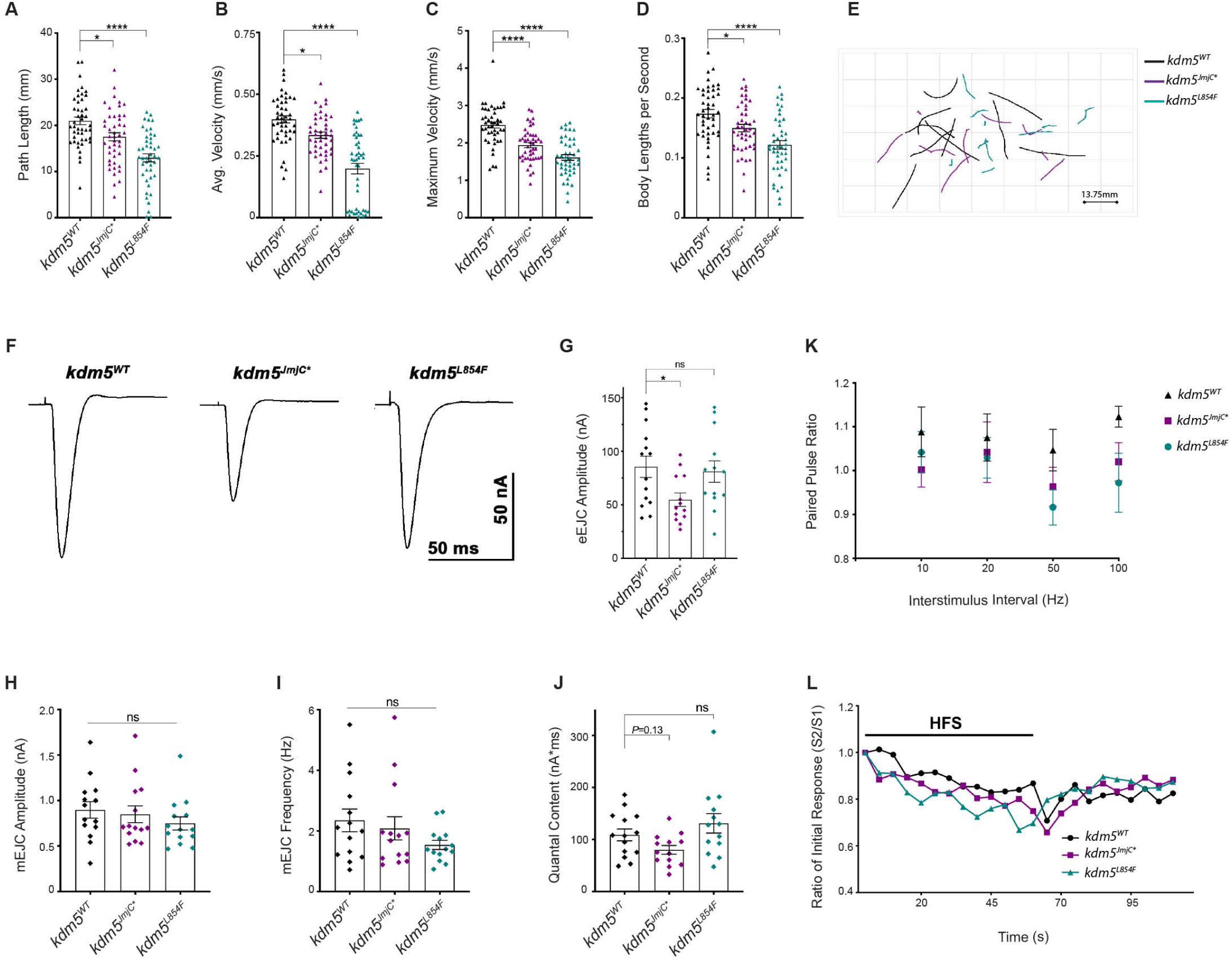
KDM5 mutations in the JmjC and C5HC2 domains affect larval locomotion. **A-E.** Larval locomotion. Movements of third instar *kdm5^WT^*, *kdm5^JmjC*^* and *kdm5^L854F^* larvae were recorded for 30 sec in an agar arena and analyzed using Worm Tracker. **A.** Quantification of traveled distance \**P*=0.0114 \*\*\*\**P*<0.0001 (one-way ANOVA Dunnett’s multiple comparisons). **B.** Quantification of average speed of movement \**P*=0.0105, \*\*\*\**P*<0.0001 (Kruskal-Wallis Dunn’s multiple comparisons). **C.** Quantification of maximum speed \*\*\*\**P*<0.0001 (one-way ANOVA Dunnett’s multiple comparisons). **D.** Quantification of the average speed normalized to larval size \**P*=0.0248 \*\*\*\**P*<0.0001 (one-way ANOVA Dunnett’s multiple comparisons). **E.** Representative path trajectories of larvae from the three strains. *kdm5^WT^* n=45, *kdm5^JmjC*^* n=45 and *kdm5^L854F^* n=45. **F.** Representative samples of evoked excitatory junctional currents (eEJCs) recorded from muscle 6 in *kdm5^WT^*, *kdm5^JmjC*^*, and *kdm5^L854F^* larvae. **G.** Quantification of eEJC amplitude. *kdm5^JmjC*^* displays a decrease in evoked current \**P*=0.0368, ns = not significant (one-way ANOVA Dunnett’s multiple comparisons). **H-I.** Quantification of miniature excitatory junctional currents amplitude (**H**) and frequency (**I**) ns = not significant (Kruskal-Wallis Dunn’s multiple comparisons). **J**. Quantification of quantal content (eEJC area (nA*ms)/mEJC area (nA*ms)), ns = not significant (Kruskal-Wallis Dunn’s multiple comparisons). **K**. Paired pulse ratio (eEJC amplitude second response/eEJC amplitude first response) across multiple interstimulus intervals. There are no differences in average ratios between *kdm5^JmjC*^* and *kdm5^WT^* at any interval (Student’s t-test). **L**. eEJC amplitudes, normalized to the first response, during high frequency stimulation (HFS) (20Hz x 60s) followed by a recovery period of stimulation (0.2 Hz x 50s). Mutant larvae show the same response as *kdm5^WT^* (Two-way ANOVA). *kdm5^WT^* n=12, *kdm5^JmjC^* n=14, and *kdm5^L854F^* n=13. Bars represent mean with SEM. In figure L error bars are not displayed to facilitate visualization.

To link the observed motor deficits to neuronal activity, we examined evoked and spontaneous synaptic transmission at the glutamatergic NMJ of larval muscle 6, which displayed similar bouton phenotypes to those observed in muscle 4 (Figure S3). Compared to *kdm5^WT^*, evoked junctional currents (eEJCs) were reduced in *kdm5^JmjC*^* but not in *kdm5^L854F^* larvae (Figure 4F-G). The impaired neurotransmission observed in *kdm5^JmjC*^* animals could be caused by defects in synaptic excitation, neurotransmitter release, or postsynaptic response. Analyses of miniature excitatory junction current (mEJC) amplitudes did not show any differences between *kdm5^JmjC*^* and *kdm5^WT^* (Figure 4H). These data suggest that the reduction in eEJCs was neither caused by a defect in the response of postsynaptic glutamatergic receptors to glutamate nor the amount of glutamate released by a single vesicle. mEJC frequency in *kdm5^JmjC*^* mutant larvae was also unaltered compared to *kdm5^WT^* indicating that there were no changes in the number of functional active release sites (Figure 4I) (Long et al., 2010). Furthermore, immunohistochemical analysis of *kdm5^JmjC*^* mutant larvae revealed no changes in the density of the presynaptic marker of active zones Bruchpilot (Brp) or it’s localization relative to glutamatergic receptors GluRIIC, in the muscle, when compared to *kdm5^WT^* controls (Figure S3F-G). We also examined quantal content (ratio of eEJC area/mEJC area), which reflects the number of vesicles released per stimulus, and observed a slight decrease in *kdm5^JmjC*^* larvae compared to *kdm5^WT^* (*P*=0.13) (Figure 4J). *kdm5^L854F^* mutant larvae did not show any differences in mEJC amplitude or frequency, quantal content or active zone density or localization (Figure 4; Figure S3). Taken together, these data indicate that loss of demethylase activity and the ID-associated mutation in the C5HC2 domain similarly impair larval locomotion but only loss of demethylase activity reduces evoked EJC amplitudes.

To evaluate whether the deficiency in neurotransmission observed in *kdm5^JmjC*^* mutants was due to differences in synaptic vesicle release probability, we analyzed the responses to paired pulse stimuli given at variable intervals (Figure 4K). The paired pulse ratio (PPR), calculated by dividing the eEJC of the second response by the first, showed no differences between *kdm5^JmjC*^* or *kdm5^L854F^* mutants and *kdm5^WT^* control larvae at any interstimulus interval. We also assessed the dynamics of synaptic vesicle recycling by recording eEJCs during a period of high frequency stimulation (HFS) (20Hz x 60s) followed by a recovery period (0.2Hz x 50s) (Long et al., 2010). Similar to controls, both mutants showed a decrease in eEJCs during HFS, when the pool of readily releasable vesicles is depleted, and an increase in eEJCs during the recovery period when the pool is replenished (Figure 4L). Together these data show that the reduction in eEJC amplitudes caused by loss of demethylase activity were not likely the result of perturbations in vesicle release probability nor endocytic recycling.

### Mutations in KDM5 that affect NMJ structure alter transcriptional programs in the VNC

As a transcriptional regulator, KDM5 likely exerts its effect on synaptic structure and function by regulating gene expression. Our results suggest that presynaptic KDM5 is important for proper synapse development (Figure 1). We therefore conducted RNA-seq using dissected VNCs where the soma of motor neurons reside. Quadruplicate RNA-seq from *kdm5^JmjC*^* and *kdm5^L854F^* mutant larvae revealed that both mutations alter gene expression relative to *kdm5^WT^*. Using an FDR cutoff of 5%, *kdm5^JmjC*^* mutant VNCs showed 1656 dysregulated genes, 858 of which were upregulated and 798 of which were downregulated (Figure 5A; Table S1). Interestingly, the genes dysregulated in the VNC of *kdm5^JmjC*^* mutants were similar to that of our previously published data from adult heads with 531 of the 685 dysregulated genes in both datasets behaving similarly (Spearman r=0.56, P<0.0001; Figure S4A-B) (Zamurrad et al., 2018). Despite this overlap, it is notable that while most ribosomal protein genes were downregulated in *kdm5^JmjC*^* adult heads leading to decreased translation, we did not observe this defect in VNC tissue (Figure S4C). These data are consistent with KDM5 regulating the expression of a subset of its target genes in a tissue- and developmental-specific manner. Comparing the VNC datasets from the two *kdm5* alleles, we found that the *kdm5^JmjC*^* allele altered expression of a greater number of genes than the *kdm5^L854F^* mutation. Using a 5% FDR cutoff, 284 genes were downregulated while 334 were upregulated in *kdm5^L854F^* mutants relative to *kdm5^WT^* (Figure 5B; Table S2). Similar to previous studies of KDM5 in *Drosophila* and other species (Chen et al., 2019; Drelon et al., 2018; Iwase et al., 2017; Mariani et al., 2016; Scandaglia et al., 2017; Zamurrad et al., 2018), the changes in gene expression were modest, averaging ~2-fold in both mutants (Figure 5A-B). It is therefore likely that KDM5 uses both its demethylase and its C5HC2 zinc finger activity to modulate the presynaptic expression of genes important for proper NMJ development and function.

**Figure 5.**
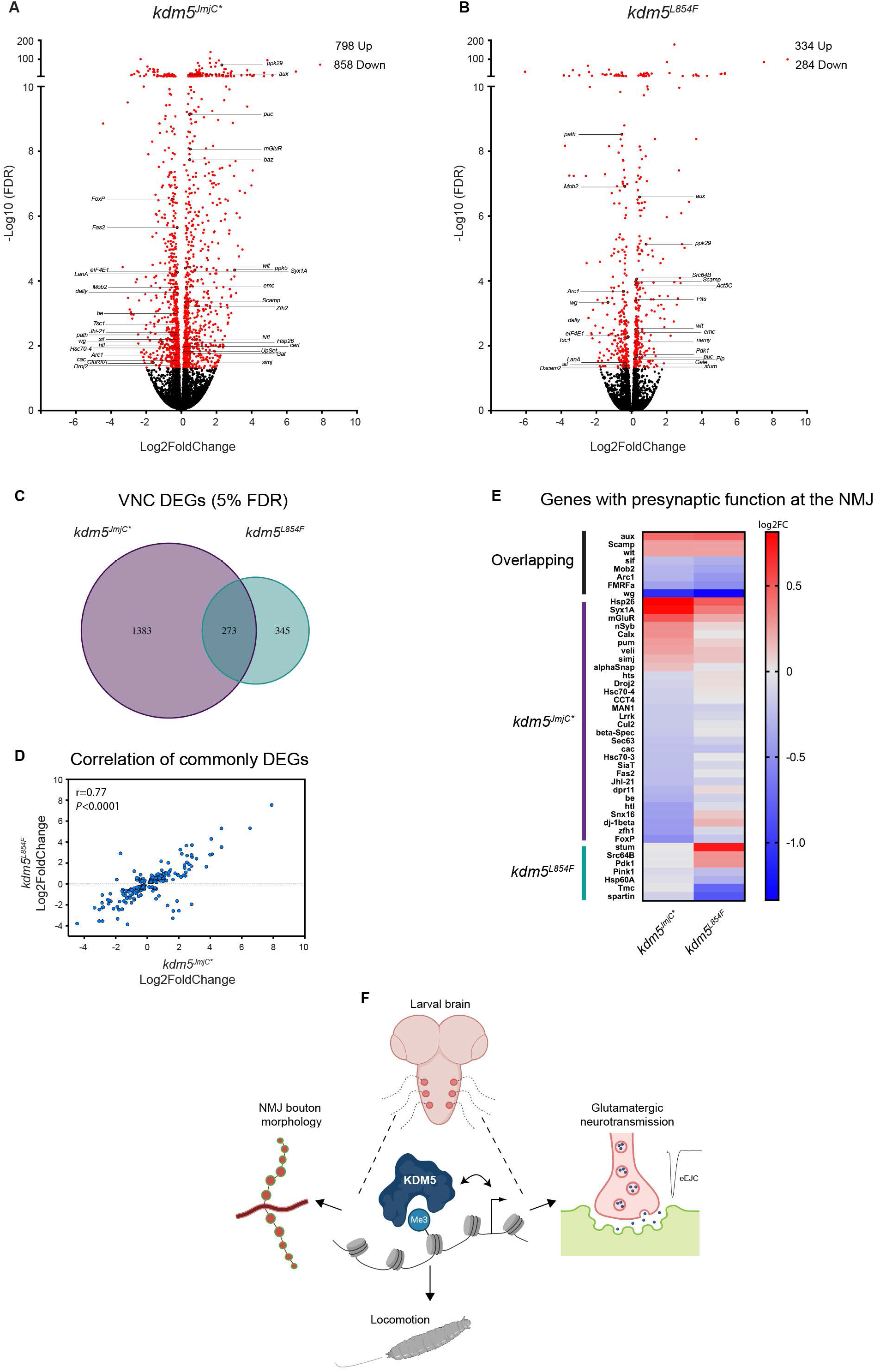
*kdm5^JmjC*^* and *kdm5^L854F^* affect KDM5-regulated gene expression programs in the larval VNC. **A-B.** Volcano plots showing dysregulated genes (5% FDR-red) in the VNC of *kdm5^JmjC*^* (**A**) and *kdm5^L854F^* (**B**). Select genes relevant to neuronal function are labeled. **C**. Venn diagram showing overlap between VNC RNA-seq data from *kdm5^JmjC*^* and *kdm5^L854F^* animals *P*=2.15e-37 (Fisher’s exact test). **D**. Correlation between changes in expression for the 273 overlapping dysregulated genes in the VNC (Spearman correlation) from *kdm5^L854F^* and *kdm5^JmjC*^*. **E**. Heatmap showing log2 fold change values of dysregulated genes with presynaptic function important for normal NMJ growth from *kdm5^L854F^* and *kdm5^JmjC*^*. The heat map is divided into sections showing genes dysregulated in both mutants and those dysregulated only in *kdm5^JmjC*^* or *kdm5^L854F^*. **F**. Model of activities of KDM5 in larval NMJs. KDM5 regulates the transcription of several genes that collectively modulate the morphology and function of the NMJ, affecting glutamatergic neurotransmission and larval locomotion.

Comparing the genes that were dysregulated in the two genotypes revealed a significant number of genes that were dysregulated by both alleles (273; *P*=2.15e-37) along with many genes that were uniquely altered in either *kdm5^JmjC*^* or *kdm5^L854F^* (Figure 5C). There was a strong correlation between commonly dysregulated genes, 243 of which behaved similarly in the two genotypes (r=0.77; 89%; Figure 5D), suggesting that these two mutations affect one or more KDM5 function(s) required at a subset of target genes. This included several genes with roles in different aspects of NMJ bouton number regulation, such as those encoding signaling molecules like Wnt/Wg pathway ligand wingless (*wg*) (Packard et al., 2002) and those involved in regulating actin cytoskeletal dynamics such as the Activity-regulated cytoskeleton associated protein 1 (*Arc1*) (Ashley et al., 2018) and the Rho family GTPase still life (*sif*) (Sone et al., 1997) (Figure 5E). Several genes important for glutamatergic synaptic function were included in the 1383 genes found to be dysregulated in *kdm5^jmjC*^* but not in *kdm5^L854F^* animals. Among these were genes encoding proteins implicated in synaptic transmission such as *cacophony* (*cac*) and *bendless* (*ben*) (Lee et al., 2014; Zhao et al., 2009) and the metabotropic glutamate receptor *(mGluR*) (Bogdanik et al., 2004; Chun-Jen Lin et al., 2011) in addition to the glutamate importer *Juvenile hormone inducible-21* (*Jhi-21*) (Ziegler et al., 2016). We also found a group of neuroprotective chaperone/stress response genes related to synaptic exocytosis and proteostasis to be dysregulated, including the Heat shock cognate genes *Hsc70-4, Hsc70-3 (Bronk et al., 2001)* and genes encoding hsp40-type proteins (*Droj2* and *Sec63*) (Santana et al., 2020) (Figure 5E). Within the 345 genes that were uniquely altered in *kdm5^L854F^*, we observed the dysregulation of genes that influence synaptic bouton size such as the Phosphoinositide-dependent kinase 1 (*Pdk1*) (Cheng et al., 2011) or that interact with, or regulate, the microtubule and actin cytoskeletons. This included genes that encode, the GTPase binding protein Spartin (*spartin*) (Nahm et al., 2013) the polymerizing GTPases of the Septin family (*pnut* and *Sep1*) (Akhmetova et al., 2018), the Rac family protein Mtl (Hakeda-Suzuki et al., 2002; Trogden and Rogers, 2015) and the kinase Src64B (Feuillette et al., 2020) (Figure 5E, Table S2). These RNA-seq data provide molecular insights into potential mechanisms by which KDM5 carries out its neuromorphological and synaptic functions.

## Discussion

Mutations in three of the four paralogs of KDM5 in humans are associated to neurodevelopmental disorders indicating a role for these proteins in neuronal development and function. We show here that KDM5 regulates the expression of genes that are collectively required for normal growth, morphology and function of the *Drosophila* larval NMJ. This synapse serves as a model for mammalian central nervous system glutamatergic synapses (Coll-Tane et al., 2019; Koh et al., 2000; Menon et al., 2013), which are altered in a number of inherited forms of cognitive impairment, including Fragile X syndrome, Angelman syndrome, and Rett syndrome (Volk et al., 2015). Specifically, our experiments demonstrate that presynaptic KDM5 promotes proper synaptic bouton number and size at the NMJ. Interestingly, while the regulation of bouton number is dependent on the enzymatic histone demethylase function and the C5HC2 zinc finger motif of KDM5, the size of the synaptic boutons depends only on the zinc finger. This suggests that the C5HC2 domain mediates key neurodevelopmental activities, which is consistent with mutations in this domain being observed in patients with ID (Jensen et al., 2005; Santos-Reboucas et al., 2011; Tzschach et al., 2006). Functionally, both *kdm5^JmJC*^* and *kdm5^L854F^* mutant larvae displayed similar locomotor defects but only *kdm5^JmjC*^* exhibited reductions in eEJC amplitudes. These overlapping, yet distinct, phenotypes of *kdm5^JmjC*^* and *kdm5^L854F^* mutants may be, at least partially, attributed to the finding that the JmjC and C5HC2 domains facilitate both shared and unique aspects of gene regulation in the larval VNC. Combined, our data lead us to propose that KDM5 uses several distinct gene-regulatory functions in NMJ glutamatergic neurons to control functional and structural neuronal characteristics.

Emerging evidence suggests that regulation of H3K4me3 is particularly important for neuronal development and function (Collins et al., 2019). Indeed, mutations in H3K4 methyltransferases, H3K4me binding proteins, and H3K4me demethylases, are found in patients with NDDs (Collins et al., 2019; Vallianatos and Iwase, 2015). Our analyses of the larval NMJ further establish the importance of KDM5-mediated transcriptional regulation and complement our previous studies showing that adult flies lacking KDM5 histone demethylase activity exhibit cognitive deficits (Zamurrad et al., 2018). Here we extend these data by demonstrating a role for the demethylase activity of KDM5 in glutamatergic synaptic transmission. Specifically, we find that evoked neurotransmission is reduced in *kdm5^JmjC*^* mutants, consistent with our transcriptomic analyses from *kdm5^JmjC*^* VNCs showing changes in the expression of genes that impact glutamatergic signaling. Because mutations in KDM5 genes typically cause modest changes to gene expression (Iwase et al., 2016; Zamurrad et al., 2018), altered expression of a relatively large number of genes in *kdm5^JmjC*^* mutant neurons could be involved in the reduced synaptic transmission. Of particular note, among the downregulated transcripts were *cac* and *ben*, which are implicated in promoting synaptic transmission and the loss of which produce a similar electrophysiological defect to *kdm5^JmjC*^* (Lee et al., 2014; Zhao et al., 2009). In addition, the expression of the gene *Hsc70-4*, involved in synaptic vesicle exocytosis (Bronk et al., 2001), was also downregulated. It is also interesting to note that the expression of the gene encoding the metabotropic glutamate receptor *mGluR* was upregulated, consistent with its role in regulating neuronal excitability by dampening neurotransmitter release (Bogdanik et al., 2004; Chun-Jen Lin et al., 2011). These data suggest that ID-associated mutations that result in the loss of histone demethylase activity are likely to cause electrophysiological changes at glutamatergic synapses. Because at least half of the known ID-associated alleles in *KDM5A*, *KDM5B* and *KDM5C* are expected to reduce protein expression or stability (Brookes et al., 2015; Lebrun et al., 2018), many patients are expected to lack histone demethylase activity. Further analyses of existing and additional ID-associated missense alleles will help clarify the precise pathways that mediate this synaptic phenotype.

In addition to exhibiting altered electrophysiology, *kdm5^JmjC*^* mutant larvae also displayed altered NMJ morphology displaying a decreased number of synaptic boutons and impaired larval locomotion. While there is often a correlation between the presence of these phenotypes, they are not necessarily directly linked (Banovic et al., 2010; Li et al., 2007; Mosca et al., 2012; Zhang et al., 2017). Indeed, changes to synaptic transmission can occur without modifying bouton morphology (Chen and Featherstone, 2011; Imlach et al., 2012) and movement deficits can arise in the absence of major changes in neurotransmission (Lembke et al., 2019). In addition, two of these defects, the reduced bouton number and the slowed larval locomotion, were shared with *kdm5^L854F^* mutants that did not show an evident decline in glutamatergic neurotransmission. One possible interpretation of our data is that the bouton number and larval movement changes that are observed in both *kdm5^JmjC*^* and *kdm5^L854F^* are functionally related, as similar correlations have been observed previously (Kashima et al., 2017). Consistent with this, RNA-seq analyses revealed a number of genes that were similarly dysregulated in the VNC of *kdm5^JmjC*^* and *kdm5^L854F^* larvae. This included diminished expression of *wingless*, which can result in reduced bouton number and slowed larval locomotion (Kim and Cho, 2020). Alternatively, the locomotor phenotype observed in *kdm5^JmjC*^* and *kdm5^L854F^* larvae may not be caused by NMJ perturbations but may instead be indirectly caused by aberrant neurotransmission in upstream motor networks. Larval locomotion is a complex behavior that relies on linking appropriate sensory inputs with a wave of peristaltic muscle contractions to mediate larval movement (Kohsaka et al., 2017). As such, it is possible that the motor deficits displayed by *kdm5^JmjC*^* and *kdm5^L854F^* mutants are caused by synaptic or morphological changes in body wall sensory neurons (Hughes and Thomas, 2007; Song et al., 2007) or defects in the premotor inhibitory GABAergic or excitatory cholinergic interneurons (Kohsaka et al., 2014). Additional studies are needed to discern the functional links between KDM5-mediated changes in bouton number, NMJ synaptic activity, and larval movement.

Our studies also emphasize the importance of non-enzymatic gene regulatory functions of KDM5 family proteins, in particular the gene-regulatory activities of the C5HC2 zinc finger motif that is affected by the L854F mutation. In addition to reduced bouton number phenotype observed in both *kdm5^JmjC*^* and *kdm5^L854F^* mutants, *kdm5^L854F^* mutant larvae also exhibited increased NMJ synaptic bouton size that correlated with changes to the microtubule cytoskeleton. Consistent with the presence of this additional structural bouton change, *kdm5^L854F^* mutant larvae displayed a number of unique gene expression defects that were not observed in *kdm5^JmjC*^* mutants. Moreover, *kdm5^L854F^* mutant CNS did not show a change to global levels of H3K4me3 as would be expected if the demethylase activity of KDM5 were the principal defect of this allele. One key group of KDM5-regulated genes whose expression is disrupted by the L854F mutation are those involved in the regulation of the microtubule dynamics, which play key structural roles and transport roles in neurons (Conde and Caceres, 2009). This observation may be important for the neuropathology of the equivalent mutation in human *KDM5C* (*KDM5C^L731F^*) as changes to presynaptic microtubule physiology are observed in other ID disorders such Fragile X syndrome (Bodaleo and Gonzalez-Billault, 2016; Lu et al., 2004). Moreover, restoring homeostasis to microtubule dynamics can attenuate morphological and functional phenotypes associated with mutations in the Fragile X gene *fmr1* (Zhang et al., 2001). Further exploration of this potential link could therefore highlight a means to attenuate the cognitive deficits for a subset of mutations associated with ID.

Because no molecular function has been attributed to C5HC2 zinc finger motifs, defining the precise molecular defect caused by changes to this domain in KDM5 family proteins will require additional analyses. This motif has previously been postulated to act as a protein-protein interaction motif and/or bind to nucleic acids (Riechmann et al., 2000; Secombe et al., 2007) Mutations in this region could therefore alter a range of functions that are critical for gene regulation, such as recruitment of KDM5 to its target genes and/or its ability to mediate activation or repression of transcription. Interestingly, the three mutations in the C5HC2 zinc finger examined do not result in the same NMJ phenotypes suggesting that they may differentially affect KDM5 transcriptional regulation on synaptic target genes. While *kdm5^L854F^* affected bouton number and size, *kdm5^Y874C^* appeared to modestly decrease bouton number, and *kdm5^R873W^* NMJs were indistinguishable from wild type. The more severe phenotypes of animals expressing the KDM5^L854F^ mutant protein are consistent with the amino acid substitution affecting one of the Zn^2+^ binding sites in the C5CH2 domain. Indeed, DynaMut predictions suggested that this amino acid variation leads to significant alterations in the dynamic and stability around the C5HC2 motif, which could cause more dramatic changes both to transcription and to NMJ morphology.

In conclusion, we show that mutations in the JmjC and C5HC2 domains of KDM5 affect the transcriptional regulation of genes that are important for synaptic structure and function. While the integrity of the JmjC demethylase domain and the C5HC2 zinc finger are each necessary to regulate the expression of a subset of genes that promote NMJ growth, these domains also carry out independent transcriptional functions in motor neurons. Specifically, the JmjC domain is required for the transcriptional regulation of genes that contribute to synaptic transmission while the C5HC2 domain regulates genes involved in cytoskeleton dynamics. Importantly, our observation that KDM5 plays critical roles at the NMJ is consistent with accumulating data linking disruption to glutamatergic synapsis to human cognitive disorders, including ID (Volk et al., 2015). Future studies will provide additional insights about the role of these domains in the recruitment of KDM5 to specific promoters.

## Materials and Methods

### Resource Availability

#### Lead Contact

Further information and requests for resources and reagents should be directed to the Lead Contact, Julie Secombe.

#### Materials Availability

All fly strains described in this manuscript are available upon request.

#### Data Availability

RNA-seq data from *kdm5^JmjC*^* and *kdm5^L854F^* mutant VNCs have been uploaded to GEO and have the accession number GSE159298. A list of differentially expressed genes (and log2 fold change) observed compared to wild type are provided in Table S1 and Table S2 (5% FDR) for *kdm5^JmjC*^* and *kdm5^L854F^*, respectively. Data from adult head RNA-seq for *kdm5^JmjC*^* compared to *kdm5^WT^* are available via accession number GSE100578.

### Care of fly strains and crosses

Unless otherwise stated, fly crosses were maintained at 25°C with 50% humidity and a 12-hour light/dark cycle. Food (per liter) contained 18g yeast, 22g molasses, 80g malt extract, 9g agar, 65 cornmeal, 2.3g methyl para-benzoic acid, 6.35ml propionic acid. The number of male and female larvae were equal across the genotypes examined. For studies comparing wild type and *kdm5* mutant larvae, animals were matched for developmental stage, not chronological age, as we previously reported (Drelon et al., 2018; Drelon et al., 2019). Wild-type wandering third instar larvae were therefore ~120 hours after egg laying (AEL) while *kdm5^140^* larvae were ~10 days old.

### Fly strains

The *kdm5^140^* null allele, *UAS*-*kdm5* transgene, and *kdm5^JmjC^*^*^ allele have been previously described (Drelon et al., 2018; Drelon et al., 2019; Navarro-Costa et al., 2016; Secombe and Eisenman, 2007; Zamurrad et al., 2018). The *UAS-shRNA* transgene to knockdown *kdm5* was generated as part of the TRiP project and has been well-validated by us and others (BL #35706) (Liu et al., 2014; Navarro-Costa et al., 2016). To generate ID alleles, we took a similar approach as our previous analyses of *kdm5^JmjC*^* (Drelon et al., 2018; Drelon et al., 2019; Navarro-Costa et al., 2016). Briefly, the 11kb genomic region encompassing *kdm5* was amplified by PCR from *w^1118^* and cloned into the pattB vector (Bischof et al., 2007) using the In-Fusion cloning system (Takara). A 3xHA tag was included in-frame at the 3’end of the *kdm5* ORF. Point mutations (L854F, R873W, Y874C) were introduced by PCR-mediated site-directed mutagenesis. Sequenced constructs were sent to BestGene for injection into FlyC31 embryos (BL #24749). Transformed flies were crossed into the *kdm5^140^* null background and homozygous stocks were established. Wild-type and demethylase dead transgenes and resulting fly strains are published (Navarro-Costa et al., 2016; Zamurrad et al., 2018). Gal4 driver fly strains used included *OK6*-*Gal4* (BL #64199), *Elav*-*Gal4* (BL#25750), *Actin*-*Gal4* (BL #3954), *da*-*Gal4* (BL #55851), *Mef2*-*Gal4* (Bl #27390), *repo*-*Gal4* (BL #7415).

### RNA-seq

Third instar larvae (pooled male and female) CNS from wild type (*kdm5^WT^*), *kdm5^JmjC*^*, and *kdm5^L854F^* were dissected Hemolymph-Like Solution (HL-3: NaCl 70mM, KCl 5mM, MgCl2 20mM, NaHCO3 10mM, trehalose 5mM, sucrose 115mM, HEPES 5mM, pH7.1) without CaCl^2^ and brain lobes were separated to enrich for VNC cells. Total RNA was isolated with Trizol (Invitrogen) and quality was assessed by Fragment Analyzer (Advanced Analytical, Ankeny, IA, USA) before sending to the Beijing Genomics Institute (BGI) for library preparation and sequencing. cDNA libraries were prepared using TruSeq Stranded mRNA Library. mRNAs were isolated from total RNA with oligo(dT) method. After mRNA fragmentation, first strand cDNA and second strand cDNA were synthesized, and cDNA fragments were purified and resolved with EB buffer for end reparation and single nucleotide A (adenine) addition. cDNA fragments were linked with adapters and those with suitable sizes were selected for PCR amplification. Libraries were sequenced on Illumina NovaSeq 6000 platform. Alignment of raw reads was performed using Bowtie2 (v2.2.5) (Langmead and Salzberg, 2012), normalized and differential expression determined with DESeq2 package (Love et al., 2014; Soneson and Delorenzi, 2013).

### Western blot

Third instar larval brains were dissected in ice-cold 1xPBS. Samples were stored in 2x NuPAGE LDS Buffer at −20°C until use. After a round of sonication and treatment with DTT, samples were subjected to SDS-PAGE and transferred to a PVDF membrane for immunoblotting. Membranes were incubated with primary antibodies at 4°C overnight, washed and incubated with secondary antibodies at room temperature for 30min. Mouse anti −histone H3, 1:2000 (Abcam #24834); rabbit anti-H3K4me3 1:2000 (Active Motif #39159); mouse anti-HA, 1:500 (Cell Signaling #2367S); rabbit anti-KDM5, 1:250 (Secombe et al., 2007); anti-gamma-tubulin, 1:1000 (4D11; Invitrogen MA1-850); and mouse anti alpha-Tubulin, 1:10000 (12G10, Developmental Studies Hybridoma Bank, University of Iowa) were used as primary antibodies. IRDye^®^ 800CW Donkey anti-Rabbit IgG (92632213) and IRDye^®^ 680RD Donkey anti-Mouse IgG (925-68072) from LI-COR were used as secondary antibodies. Western blots were quantified using LI-COR Image Studio software.

### Immunostaining

Third instar larvae were dissected in ice-cold 1xPBS and fillets were fixed in 4%PFA in PBS at room temperature for 30min. For Futsch staining, fillets were fixed in ice-cold methanol for 10min at −20°C. Samples were washed in 1xPBS for 10 min followed by two washes in 1xPBST (PBS + 0.1% Triton) for 10min each. Fillets were transferred to 0.5μl tubes for primary antibody incubation overnight at 4°C with agitation. After three 15 min washes in 1xPBST, samples were incubated in secondary antibodies for 1 h at room temperature. Samples were then washed three times for 10 min each and mounted with Fluoromount-G™ DAPI (SouthernBiotech) or SlowFade™ Glass Antifade Mountant (Invitrogen). Mouse anti-Dlg (4F3) 1:1000, anti-Futsch (22C10) 1:100, and anti Brp (nc82) 1:100 were obtained from Developmental Studies Hybridoma Bank (DSHB, University of Iowa). Rabbit Anti-GluRIIC (1:5000) was a kind gift from Aaron DiAntonio. Alexa 647-conjugated goat anti-HRP was used at 1:200 (Jackson ImmunoResearch Laboratories, Inc., West Grove, PA). Goat anti-mouse Alexa 488 (Invitrogen) was used at 1:400 while anti-mouse TRITC and anti-rabbit FITC (Jackson ImmunoResearch Laboratories, Inc., West Grove, PA) were used at 1:400.

### Electrophysiology

Third instar larvae were fillet dissected in ice cold HL-3.1 (Feng et al., 2004) containing 0.25 mM Ca^2+^. Larvae were glued (Vetbond Tissue Adhesive, World Precision Instruments) to Sylgard-coated coverslips, the VNC was cut out of the animal, and the HL-3.1 was replaced with room temperature HL-3.1 containing 1.0 mM Ca^2+^ for recordings. Muscle 6 in segments A3 or A4 was voltage clamped at −60 mV using an Axoclamp 900A amplier (Molecular Devices). Clamp and recording electrodes with 10-20 MΩ of resistance were filled with 3 mM KCl. Supra threshold stimuli were delivered via stimulating electrodes filled with bath solution using a Grass S88 stimulator with a SIU5 isolation unit (Grass Technologies). Evoked EJCs were measured after stimulating segmental nerves at 0.2 Hz for 50 s to establish baseline responses. Endocytosis was assessed by stimulating by 20 Hz for 60 s. Recovery of the cycling pool of vesicles was assessed by stimulating at 0.2 Hz for 50 s. Paired pulse amplitudes were measured after delivering two 10 Hz, two 20 Hz, two 50 Hz, and two 100 Hz pulses, each of which were separated by a 20 s intertrial interval. Recordings were digitized with a Digidata 1443 digitizer (Molecular Devices). Quantal content was calculated by dividing the eEJC area (nA × ms) by the mEJC area (nA × ms) for each animal. 180 s of spontaneous activity was used to quantify mEJC frequency and amplitude. An equal number of recordings from controls and experimental animals were obtained each day. Data were analyzed in pClamp (v11.0, Molecular Devices).

### Larval movement assays

Third instar larvae were placed on Petri dishes containing 1.6% agar in double distilled water. Larvae were allowed to crawl on the agar dish for one minute to remove excess food and then transferred to the crawling arena, which also consisted of 1.6% agar. Larvae were allowed to acclimate for one minute then larval locomotion was recorded using a Cannon EOS M50 camera at 29.97 frames per second for 30 seconds. Five larvae were recorded per video. Videos were opened in ImageJ (NIH) and 899 frames were analyzed with the wrMTrck plugin (Jesper S. Pedersen).

### Developmental timing quantification

Female and male flies were placed in a vial and allowed to lay eggs for 24 hours. Starting at day 5 after egg laying the number of animals that had pupariated were scored twice per day.

### Image acquisition and processing

Images of NMJs from muscles 4 and 6 of abdominal segment 3 were acquired using Leica Sp8 Confocal Microscope, 63x/1.4NA and Zeiss Axioimager Z1 Apotome. Quantification of muscle area and type Ib bouton number and area were performed in the Dlg channel using ImageJ. The number of boutons was normalized to muscle surface area similar to previous studies (Banovic et al., 2010; Nechipurenko and Broihier, 2012). The Threshold and Measure commands were used to quantify bouton surface areas, similar to previous studies (Wagner et al., 2015). Adult fly images were obtained using a stereomicroscope Carl Zeiss Stereo Discovery V12 with 14X magnification and captured using AxioVision Release 4.8 software. All images were processed and assembled into figure format using Adobe Illustrator 2020. Images of adult head and ventral nerve cord and icons used in the model were created with BioRender.com.

Colocalization of Brp and GluRIIC was quantified using Fiji (NIH ImageJ) by outlining the NMJ in a max-projected image. Pearson’s R values were obtained for each image after splitting the channels using the Coloc2 plugin.

### *In silico* mutation analysis

Normal mode analysis was used to predict the thermodynamic changes upon mutations in the human KDM5A structure (PDB:5CEH) via the DynaMut server (Rodrigues et al., 2018). Files generated by DynaMut were visualized using PyMol (The PyMOL Molecular Graphics System, Version 2.4 Schrödinger, LLC). UCSF’s Chimera (Pettersen et al., 2004) was used to modify the KDM5A structure.

### Statistical analyses

All experiments were performed in biological triplicate (minimum) and numbers (N) are provided for each experiment either in the figure or the legend. Fisher’s exact test was performed using R. Student’s *t*-test, one-way ANOVA, Kruskal-Wallis tests and Two-way ANOVA were calculated using GraphPad Prism (v 8.4).

## Acknowledgements

We would like to thank members of the Secombe and Liebl labs for insights at all stages of this project. We also thank Jacqueline Tobin for her help as a summer student during this project. We are grateful for the GluRIIC antibody form Dr. Aaron DiAntonio lab, fly strains from the Bloomington *Drosophila* Stock Center (NIH P400D018537) and antibodies from the Developmental Studies Hybridoma bank, created by the NICHD of the NIH and maintained at The University of Iowa. We also thank the NIH for funding to J.S. (R01 GM112783) and F.L. (1R15NS101608-01A1), the shared instrument grant 1S10OD023591-01, and the Einstein Cancer Center Support Grant P30 CA013330.

## Author Contribution

Conceptualization, J.S., H.B., F.L.; Methodology H.B.; Investigation, H.B., S.Z., E.H., F.L. and J.S.; Writing – original draft, H.B and J.S., Writing – Reviewing and Editing, H.B., J.S., F.L., and E.H‥; Funding acquisition, J.S. and F.L, Supervision, J.S. and F.L.

## Competing interests

The authors declare no competing interests.

## Funding

This work was supported by the National Institutes of Health [R01GM112783 to J.S; 1R15NS101608-01A1 to F.L.]. The content is solely the responsibility of the authors and does not necessarily represent the official views of the NIH.

---

**Supplementary Figure 1.**
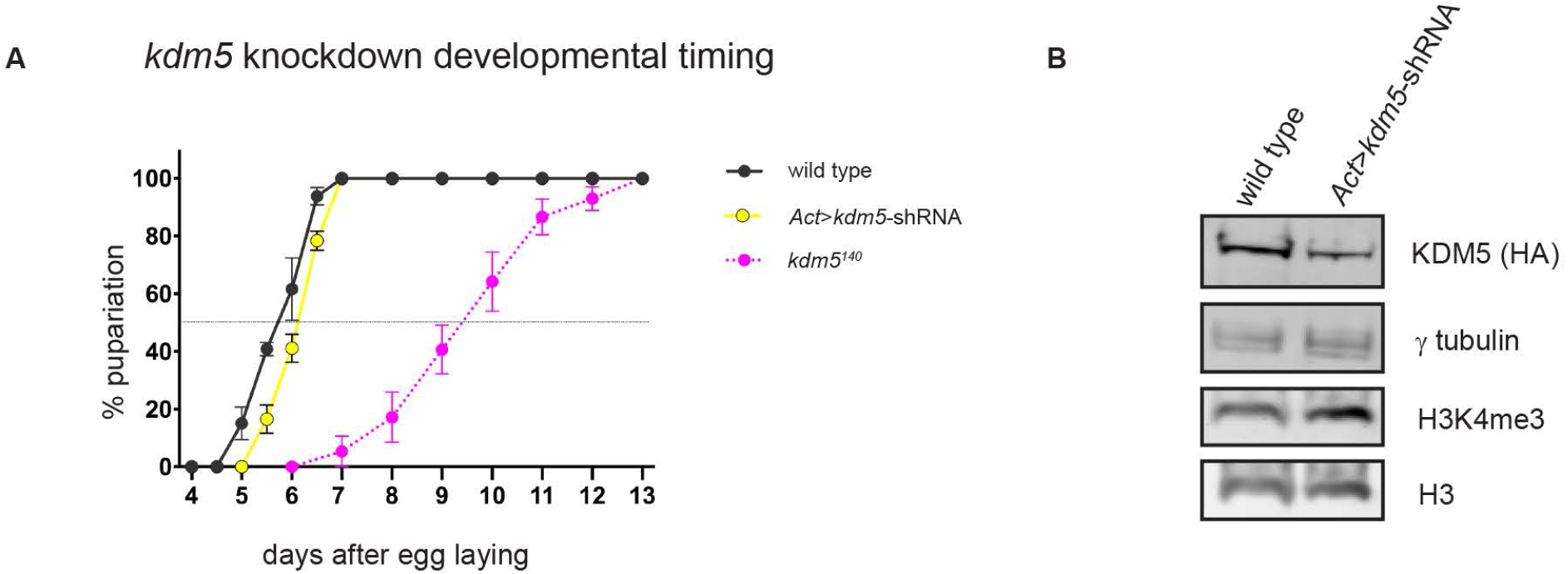
Kdm5 ubiquitous knockdown affects KDM5 demethylase activity without grossly affecting developmental timing. **A**. Developmental timing of the control strain *w^1118^*, the null mutant *kdm5^140^*, and the *kdm5* knockdown *Act*>*kdm5*-shRNA determining the day when 50% of the animals reach pupariation. Total pupae quantified in three different experiments. *w^1118^* (n=87), *kdm5^140^* (n=49), *Act*>*kdm5*-shRNA (n=79). **B**. Western blot using dissected larval CNS material from *w^1118^* and *kdm5* knockdown *Act*>*kdm5-*shRNA showing expression of HA-tagged KDM5, γ tubulin, H3K4me3 and histone H3.

**Supplementary Figure 2.**
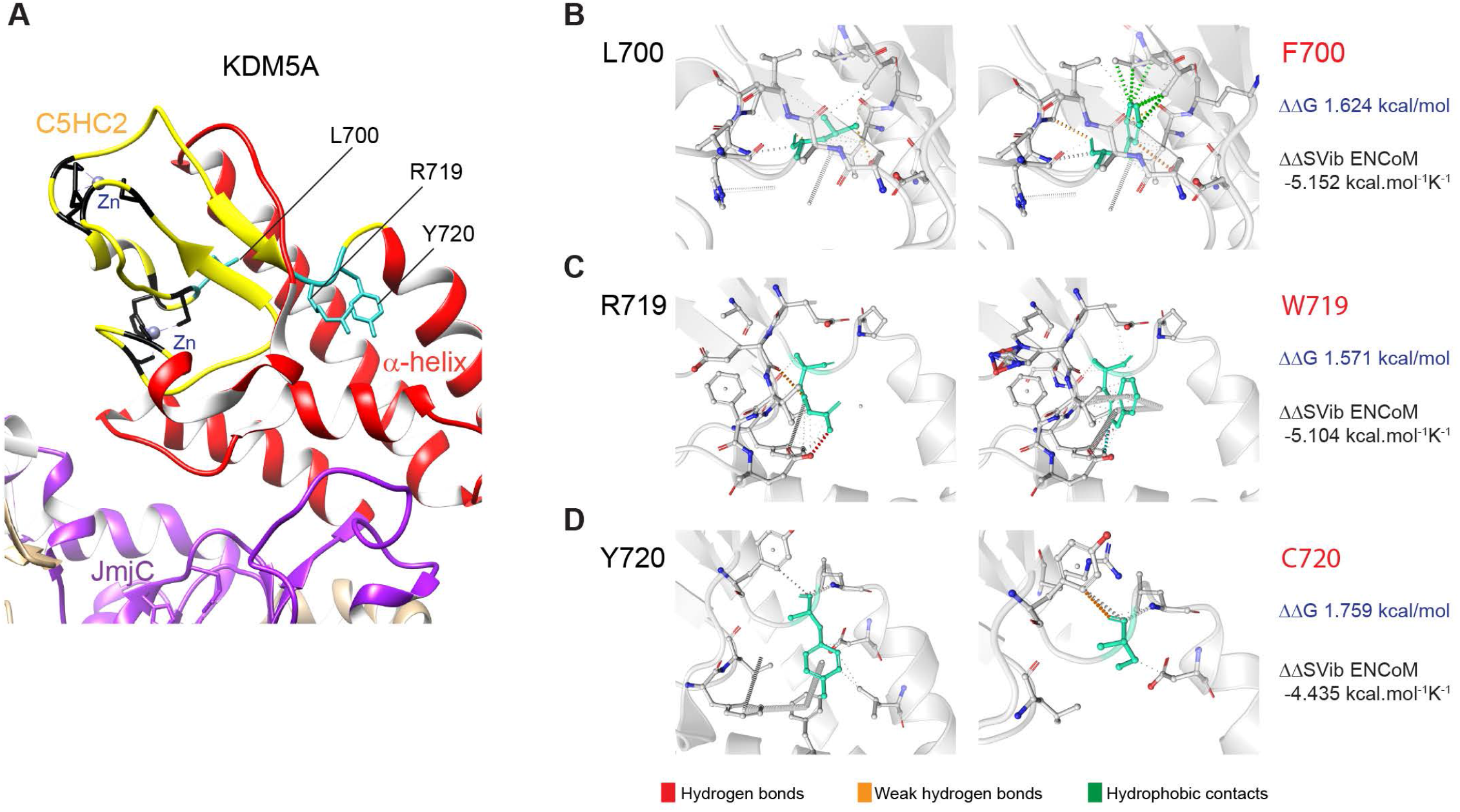
ID mutations in the C5CH2 domain of KDM5 may lead to changes in thermodynamic stabilization and molecule flexibility of the protein. **A.** Crystal structure of human KDM5A (PDB:5CEH) showing the amino acids altered by ID-associated mutations in the C5HC2 domain that were analyzed in this study. **B-D.** Predicted intramolecular interactions of the wild-type L700, R719, Y720 and mutated residues F700, W719 and C720 with the surrounding residue environment (corresponding amino acids in *Drosophila*: L854F, R873W, Y874C). The change in Gibbs free energy (ΔΔG) calculated by DynaMut predicts stabilization of the structure for the three missense mutations (ΔΔG>0). Calculations of the vibrational entropy energy (ΔΔSVib) predict decrease of molecule flexibility for the three mutations (ΔΔSVib<0).

**Supplementary Figure 3.**
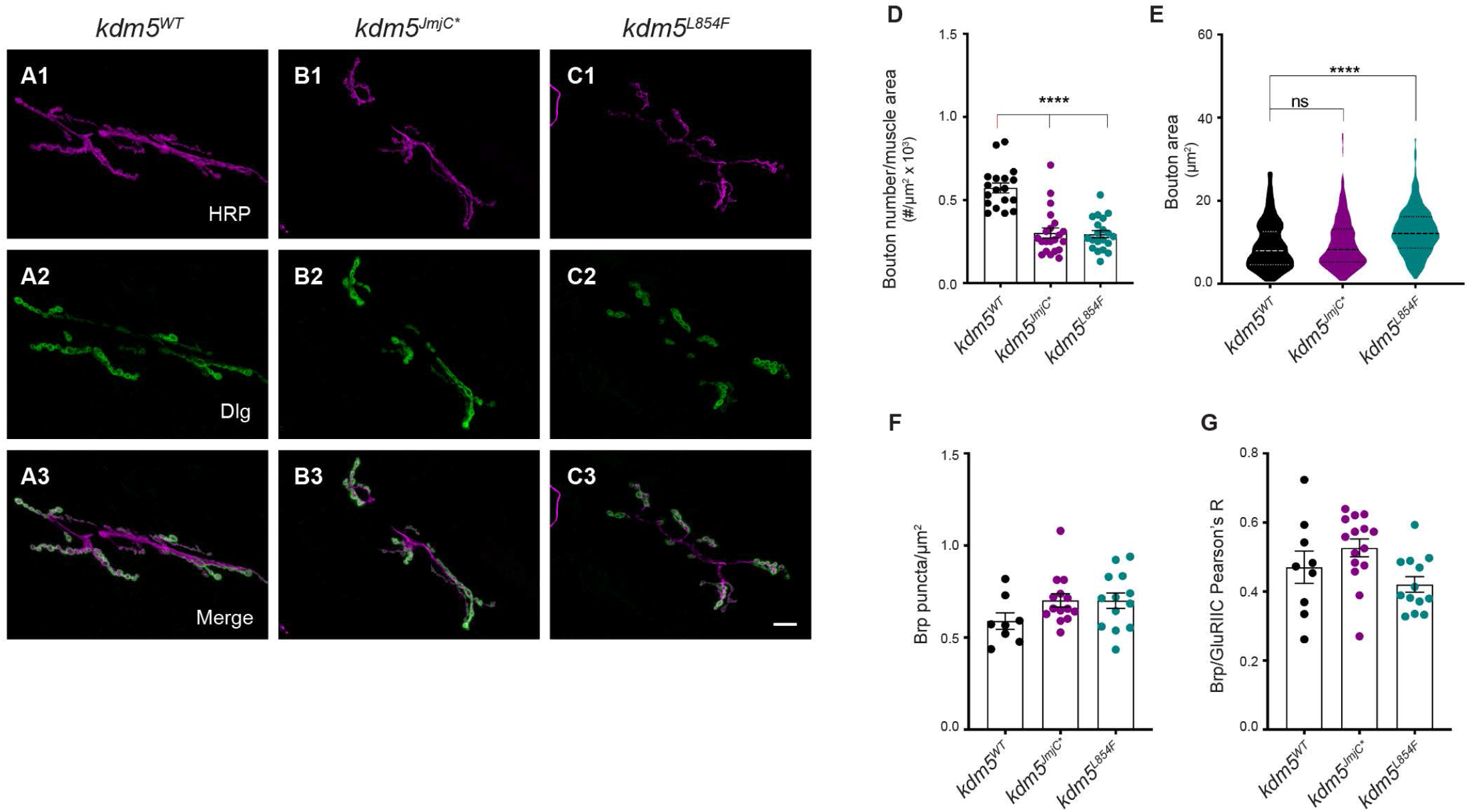
*kdm5^JmjC*^* and *kdm5^L854F^* mutants display altered 6/7 NMJ morphology but no changes in and the number or localization of active zones. **A1-C3.** 6/7 NMJ morphology at A3 of third instar larvae labeled with HRP (magenta) and Dlg (green). Compared to *kdm5^WT^* (**A1-3**), *kdm5^JmjC*^* (**B1-3**) and *kdm5^L854F^* (**C1-3**) larvae display a reduction in the number of type Ib synaptic boutons. Scale bar, 20μm. **D**. Quantification of type Ib bouton number normalized to muscle surface area for *kdm5^WT^*, *kdm5^JmjC*^* and *kdm5^L854F^*. \*\*\*\**P*<0.0001 (Kruskal-Wallis Dunn’s multiple comparisons). *kdm5^WT^* n=18, *kdm5^JmjC*^* n=21, *kdm5^L854F^* n=20. **E**. Quantification of type Ib bouton size. Violin plots show the frequency distribution of bouton surface area indicating the median and quartiles. \*\*\*\**P*<0.0001 ns = not significant (Kruskal-Wallis Dunn’s multiple comparisons). *kdm5^WT^* n=306, *kdm5^JmjC*^* n=195, *kdm5^L854F^* n=160. **F**. Quantification of Brp density calculated as number of fluorescent puncta divided by total NMJ area. ns = not significant (Kruskal-Wallis Dunn’s multiple comparisons). *kdm5^WT^* n=8, *kdm5^JmjC*^* n=14, *kdm5^L854F^* n=13. **G**. Quantification of Brp/GluRIIC apposition. ns = not significant (one-way ANOVA Dunnett’s multiple comparisons). *kdm5^WT^* n=9, *kdm5^JmjC*^* n=15, *kdm5^L854F^* n=13. Bars represent mean with SEM.

**Supplementary Figure 4.**
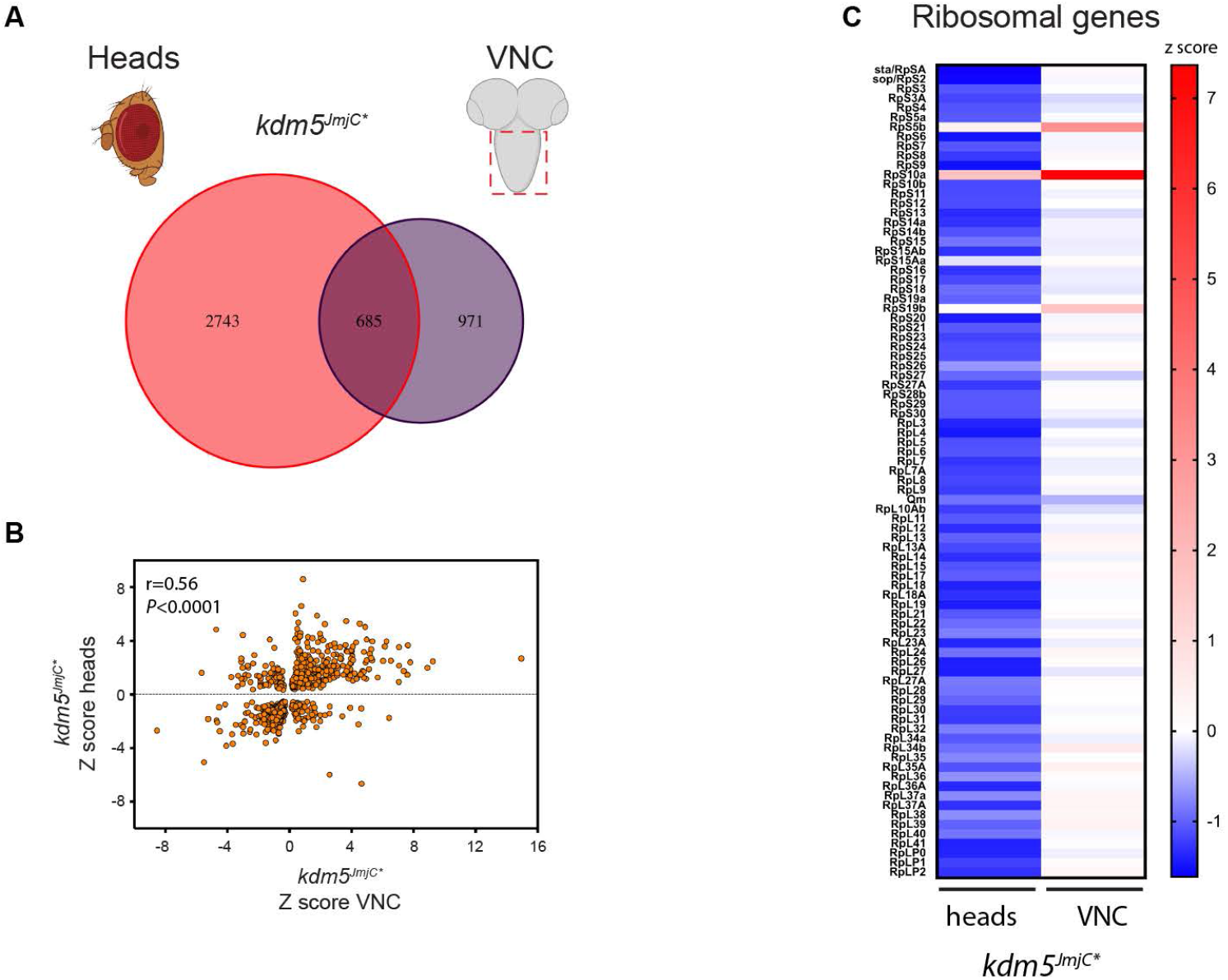
Comparison of transcriptional profiles of adult head and larval VNC mutants. **A**. Venn diagram showing overlap between VNC RNA-seq data and adult heads RNA-seq data (GSE100578) for *kdm5^JmjC*^ P*=1.8e-79 (Fisher’s exact test). **B**. Correlation between expression changes for overlapping genes in C using z scores (Spearman correlation). **C**. Heatmap of RNA-seq expression z-scores calculated for all ribosomal genes from *kdm5^JmjC*^* VNC and heads samples.

## Notes

### Competing Interest Statement

The authors have declared no competing interest.

